# Mapping the invisible chromatin transactions of prophase chromosome remodelling

**DOI:** 10.1101/2021.06.21.449273

**Authors:** Itaru Samejima, Christos Spanos, Kumiko Samejima, Juri Rappsilber, Georg Kustatscher, William C. Earnshaw

**Author notes:** Communicating Authors.

## Abstract

We have used a combination of chemical genetics, chromatin proteomics and imaging to map the earliest chromatin transactions during vertebrate cell entry into mitosis. Chicken DT40 CDK1^as^ cells undergo synchronous mitotic entry within 15 minutes following release from a 1NM-PP1-induced arrest in late G_2_. In addition to changes in chromatin association with nuclear pores and the nuclear envelope, earliest prophase is dominated by changes in the association of ribonucleoproteins with chromatin, particularly in the nucleolus, where pre-rRNA processing factors leave chromatin significantly before RNA polymerase I. Nuclear envelope barrier function is lost early in prophase and cytoplasmic proteins begin to accumulate on the chromatin. As a result, outer kinetochore assembly appears complete by nuclear envelope breakdown (NEBD). Most interphase chromatin proteins remain associated with chromatin until NEBD, after which their levels drop sharply. An interactive proteomic map of chromatin transactions during mitotic entry is available as a resource at https://mitoChEP.bio.ed.ac.uk.

## INTRODUCTION

Mitotic chromosomes and interphase chromatin differ dramatically in appearance and composition. This reflects distinct functional requirements (e.g., regulated gene expression and maintenance versus chromosome segregation) involving different organization of the chromatin fiber. Interphase nuclei are hierarchical ensembles of local chromatin folding (e.g., TADs) and functional segregation into compartments (Dekker and Mirny, 2016). Mitotic chromosomes are linear arrays of loops organized by an interplay between nuclear condensin II and cytoplasmic condensin I (Gibcus et al., 2018; Naumova et al., 2013). Currently, little is known about how interphase chromatin structures are disassembled and the chromatin proteome is reorganised during formation of the rod-liked mitotic chromosomes (Hirano, 2015; Paulson et al., 2021; Takahashi and Hirota, 2019; Zhou and Heald, 2020).

Mitotic entry is driven by a kinase/phosphatase network in which activating and inhibitory factors shuttle between the cytoplasm and nucleus (Hagting et al., 1999). Physiologically, mitosis begins with Cdk1-cyclin B1 activation on centrosomes (Jackman et al., 2003) followed by Cdk1-cyclin A2-dependent migration of cyclin B1 into the nucleus (Hegarat et al., 2020). Use of a FRET reporter found that the earliest visible consequence of Cdk1 activation in HeLa cells was cell rounding (Gavet and Pines, 2010).

Chromosome condensation is the key cytological landmark that classically defines the beginning of prophase (Flemming, 1882). Because interphase chromatin reorganization into individual mitotic chromosomes is extremely subtle at first, it is difficult to define exactly when the process begins. Thus, the early events of mitotic chromosome formation remain relatively inaccessible.

We have used chemical genetics to map early prophase events. Pioneering work of Shokat recognized that some kinases retain catalytic activity following replacement of a bulky “gatekeeper” residue near the ATP-binding pocket with a smaller residue (Bishop et al., 1998; Bishop et al., 2000). This allows the inhibitor 1NM-PP1 to dock, preventing ATP-binding and inactivating the kinase. 1NM-PP1 has the advantages that −1- it is highly specific for the engineered kinase, and −2- it can be washed out relatively quickly and efficiently (Bishop et al., 2001; Gibcus et al., 2018; Samejima et al., 2018).

CDK1 is essential for mitotic entry (Nurse, 1990; Santamaria et al., 2007). Exploiting the evolutionarily conserved nature of CDK1 (Lee and Nurse, 1987), we made a sub-line of chicken DT40 cells whose cell cycle is driven by *Xenopus* CDK1^as^. *Xenopus* CDK1^F80G^ is 1NM-PP1-sensitive (CDK1^as^) and retains sufficient activity to drive the growth of cultured cells (Hochegger et al., 2007). 1NM-PP1 treatment causes these DT40 cells to accumulate in late G_2_, and they enter mitosis within a few minutes of inhibitor wash-out (Gibcus et al., 2018; Samejima et al., 2018). We previously exploited the synchronous mitotic entry of DT40 CDK1^as^ cells to study changes in the folding of the chromatin fiber during mitotic chromosome formation (Gibcus et al., 2018).

Here, we present a chromatin map of the earliest events of mitotic entry starting well before any visible sign of mitotic chromosome formation. We find that the earliest prophase chromatin changes occur at nuclear pores, on the inner surface of the nuclear envelope, and most strikingly, in the nucleolus. There, proteins involved in rRNA processing move away from the chromatin, leaving behind the RNA polymerase I (RNAPI) machinery. Our work defines successive waves of chromatin proteome remodeling that accompany nuclear disassembly and mitotic chromosome formation.

## RESULTS

### Proteomic profiling of chromatin during mitotic entry

To study chromatin proteome dynamics during mitotic entry we used a chemical-genetic system that allows us to obtain highly synchronous populations of chicken DT40 cells entering prophase (Gibcus et al., 2018). The protocol is shown in Figure 1A.

**Figure 1.**
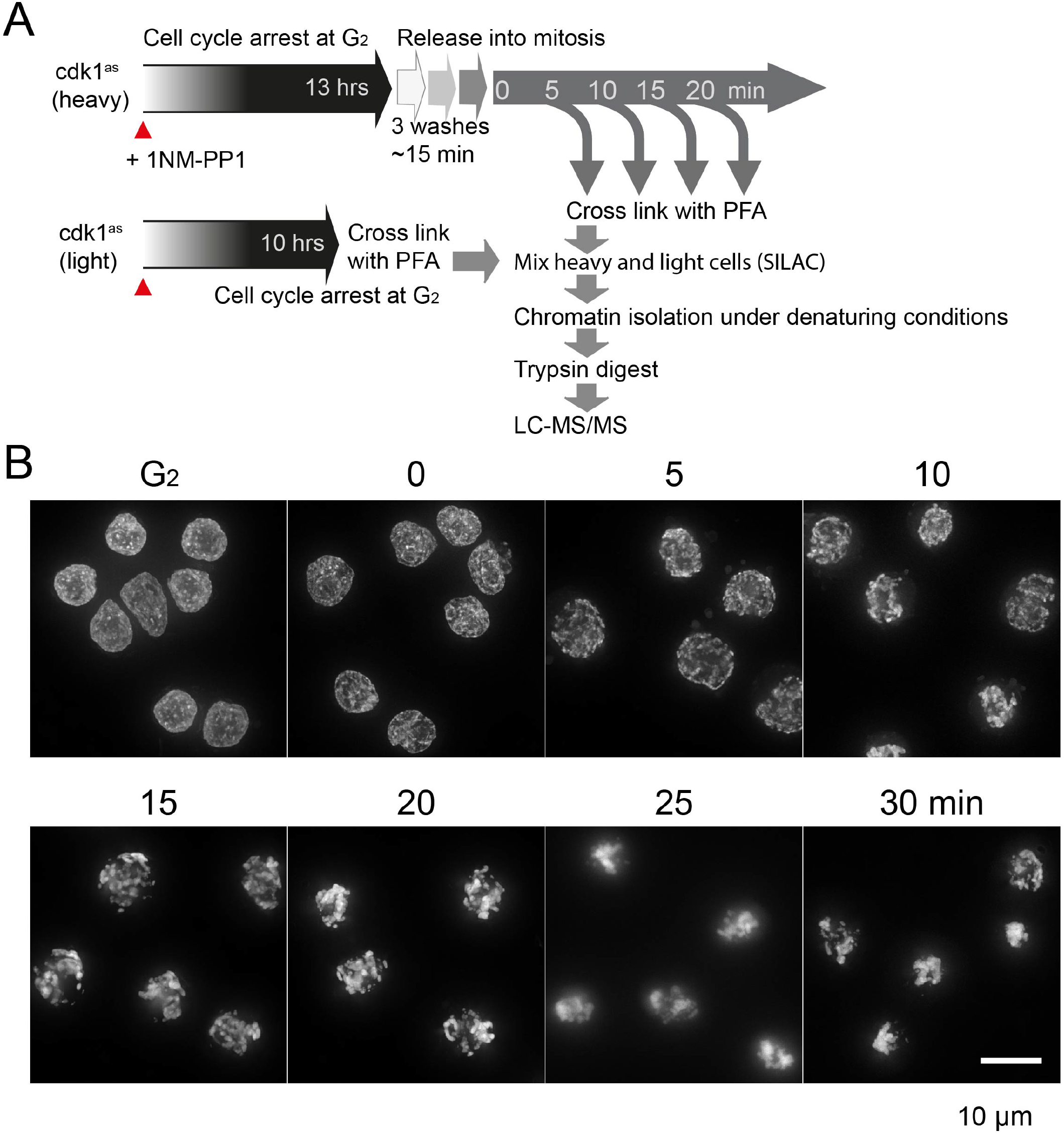
Proteomic profiling of synchronous mitotic entry. (A) Workflow of time-resolved mitotic chromatin proteomics. DT40 CDK1^as^ cells arrested in G_2_ by 1NM-PP1 were released into mitosis by washout of the drug. Crosslinked cells were processed by ChEP and by LC MS/MS (Kustatscher et al., 2014b). (B) Synchronous mitotic entry of DT40 CDK1^as^ cells. Cells were fixed with 4% formaldehyde and stained with Hoechst at the indicated times after 1NM-PP1 washout. Images are projections of z stacks. Scale bar, 10 μm.

In brief, chicken DT40 cultures whose cell cycle is driven by analogue sensitive *Xenopus* CDK1^as^ were arrested in late G_2_ with the ATP analog 1NM-PP1. 1NM-PP1 washout, involving three centrifugations, activates the kinase, triggering rapid mitotic entry (Figure 1B). In our experiments, T = 0 corresponds to the completion of the centrifugations, ~15 minutes after the first drop in 1NM-PP1 levels. We do not start our timelines with the first 1NM-PP1 wash-out because on different days slight differences in sample handling can cause a variation of 1-2 minutes in the washing time. At the end of our analysis (T= 25 minutes) the cells are fully established in prometaphase.

The micrographs of Figure1B highlight the synchronous mitotic entry following Cdk1 activation. The chromatin distribution is altered already by 5 minutes as the vast majority (>90%) of cells enter prophase. Prophase chromosome formation appears complete by 10 minutes. By 15 minutes >90% of the cells are in prometaphase.

These synchronous mitotic populations offer two important advantages. Firstly, given the rapid and synchronous mitotic entry, we can study events of prophase that occur before there is any visible evidence of mitotic chromosome condensation. Secondly, the high degree of synchrony allows biochemical analysis of events that could previously only be studied by live-cell microscopy. Our analysis uses Chromatin Enrichment for Proteomics (ChEP), which, like chromatin immunoprecipitation (ChIP), detects the susceptibility of proteins to be formaldehyde crosslinked to DNA (Kustatscher et al., 2014a; Kustatscher et al., 2014b). To measure quantitative changes with high accuracy by LC-MS/MS, isotope-labelled samples from each time point were combined with a light G_2_/M-arrested reference population prior to chromatin fractionation and analysis (Figure 1A).

### Extensive remodeling of the chromosome proteome during prophase

Our analysis identified 2,592 proteins at all time points in two biological replicates with strong reproducibility (Figure S1, Table S1). These proteins show diverse kinetic profiles (Figure 2A; Figures S2 S3). To summarise, during mitotic entry 1312 (~50%) of the proteins were depleted in chromatin, while 524 (20%) accumulated. 756 proteins (29%) exhibited only minor changes in chromatin association. Early prophase was dominated by proteins moving away from chromatin but accumulation was already evident for some, including some cytoplasmic proteins, by T= 5 minutes.

**Figure 2.**
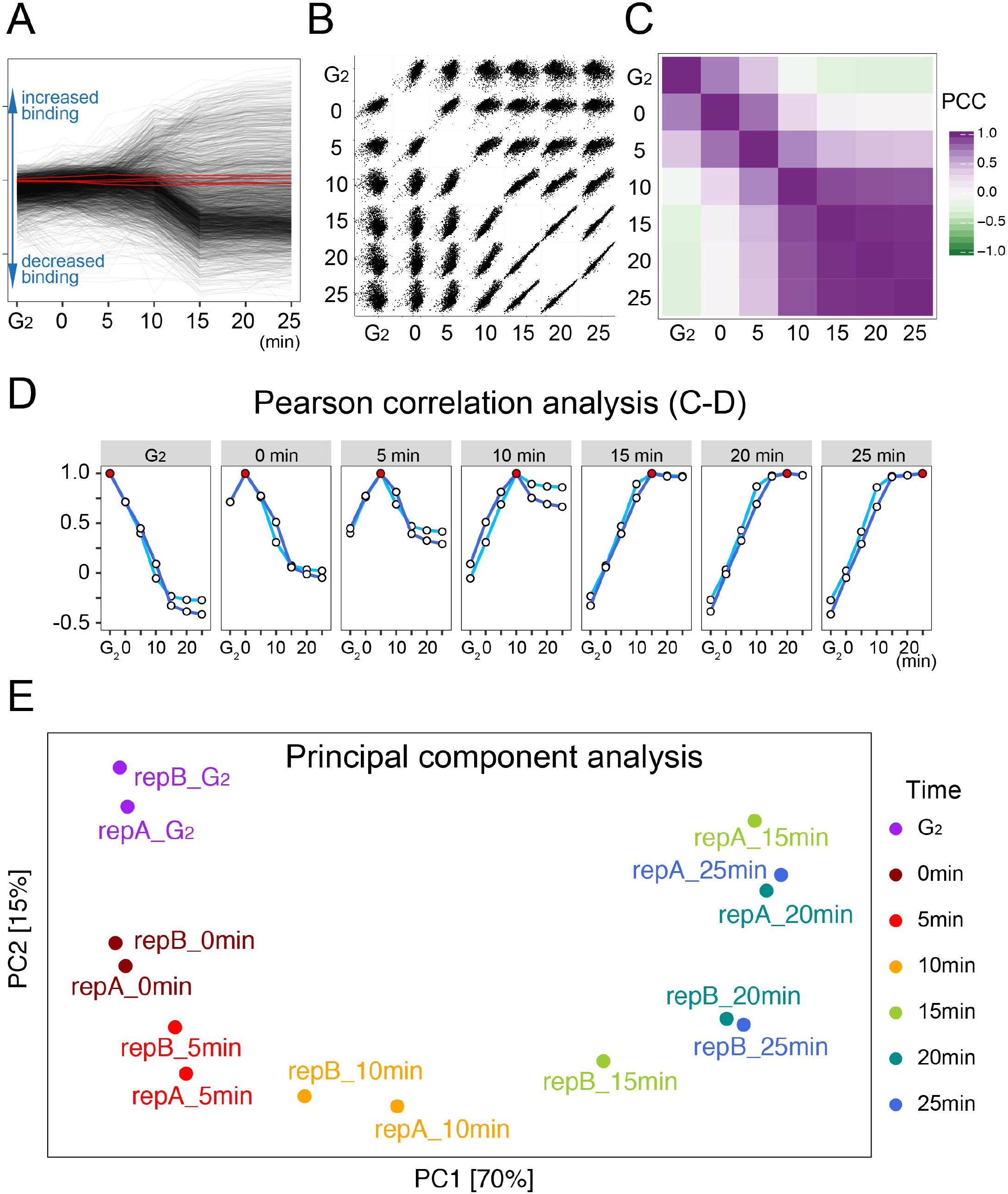
Extensive remodeling of the chromosome proteome during prophase. (A) General overview of chromatin protein changes during mitotic entry. Red lines correspond to the core histones. (B) Scatter plots comparing SILAC ratios of all proteins between time points. (C) Heat map illustrating Pearson correlation coefficients (PCC) between time points. (D) Line plots comparing the PCC between a designated time point (red dot) and all other time points. Data from two replicates are shown. (E) Principal component analysis of the proteomic time course samples.

Early prophase is a time of widespread dramatic remodeling of the chromatin proteome that begins shortly after 1NM-PP1 washout. This is evident from the fact that protein levels of early time points (G_2_ to 5 min) are correlated poorly with each other or any other time point (Figure 2). These dynamic changes cease at the time of NEBD, and protein fold-changes are highly correlated across the later time points (≥15 minutes, Figure 2B-D). Ten minutes is a clear transition point, with characteristics of both prophase and prometaphase (Figure 1B). This conclusion was confirmed using principal component analysis (Figure 2E).

### Loss of nuclear envelope barrier function during early prophase

To understand the reason for this transition at 10 minutes in DT40 CDK1^as^ cells we monitored the behavior of two fluorescently labelled reporter constructs during mitotic entry (Figure 3). 3xGFP with a nuclear localization signal (3xGFP-NLS) was used to detect nuclear-cytoplasmic mixing in live cells. Lamin B1 (LMNB1) was labeled by N-terminal Halo tag knock-in to stain the lamin network in fixed cells.

**Figure 3.**
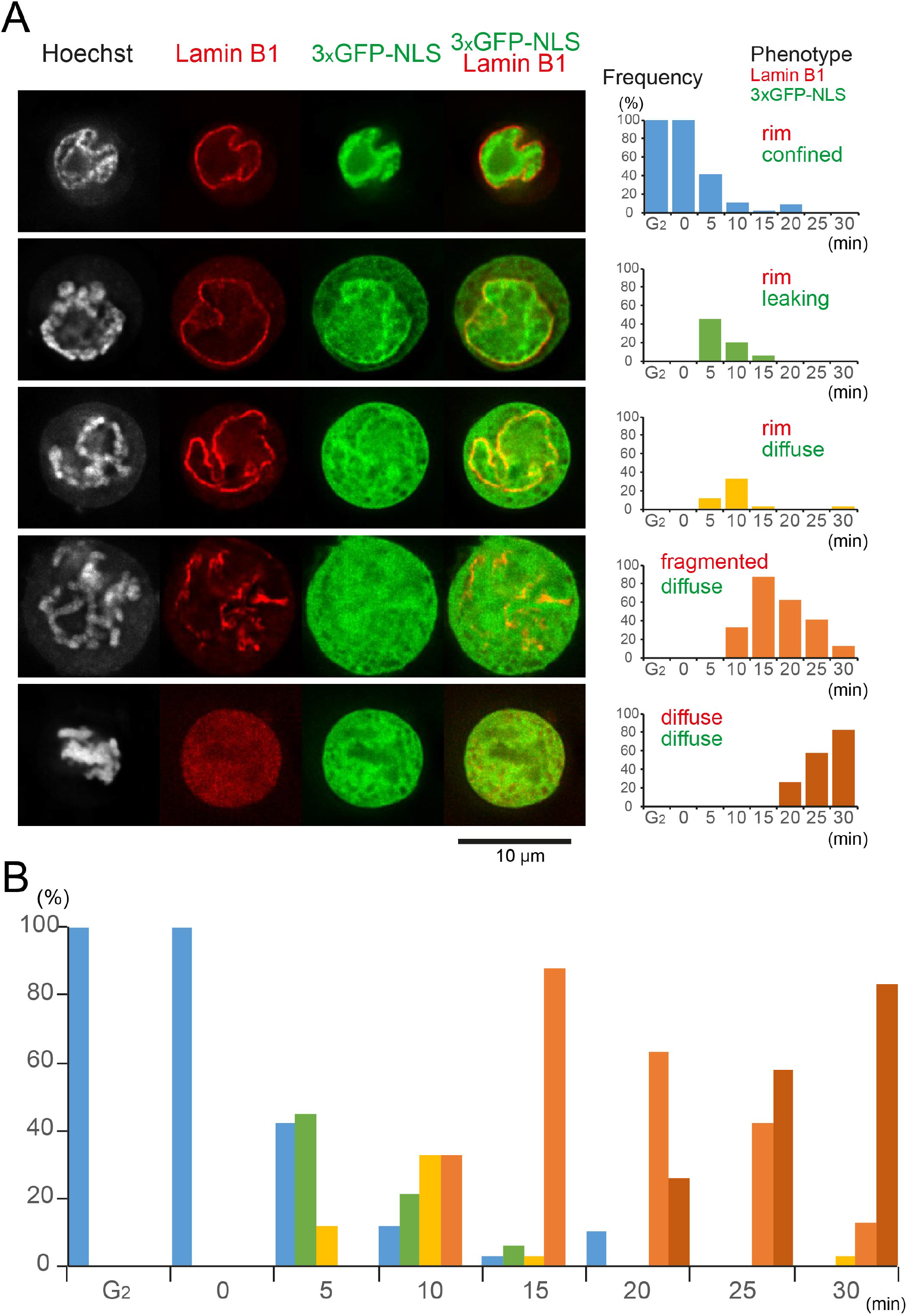
Nuclear envelope partially loses barrier function before complete breakdown. (A) Cell populations at different mitotic time points were classified according to localization patterns of lamin B1 and GFP-NLS. Bar graphs show occurrence of each phenotype at each time point. Images show single z slices. (B) Distribution of each class at each time point.

The nuclear lamina localised to the nuclear rim in all cells from G_2_ through 5 minutes in our time course (Figure 3A). This persisted in > 60% of cells at 10 minutes but the lamina was obviously fragmented by 15 minutes in most cells. From 20 min onwards Lamin B1 was largely diffuse throughout the cell.

The behavior of the 3xGFP-NLS reporter was uncoupled from that of the lamina, with leakage into the cytoplasm readily apparent at 5 minutes despite an apparently intact lamina, (Figure 3A, 2^nd^ row, green bars). By 10 minutes, the 3xGFP-NLS signal was evenly dispersed throughout the cell in >60% of cells, even though the lamina was fragmented in only 33% of cells. Thus, the 10-minute time point represents a mixture of prophase and prometaphase cells, as suggested by the correlation analyses of Figure 2. During prometaphase, both Lamin B1 and the 3xGFP-NLS reporter are diffuse throughout the cell (Figure 3A).

An integrated plot for the entire time course (Figure 3B) shows clearly that cytoplasmic factors can access and function in the chromatin much earlier than previously thought. The changes in nuclear envelope permeability are accompanied by early changes in the chromatin association of nuclear pore and inner nuclear envelope proteins (see below).

### Classifying patterns of chromosomal protein behavior during mitotic entry

To capture large trends in chromatin proteome remodeling during mitotic entry we grouped proteins by *k*-means clustering. Dividing the proteomic time course into six clusters (*k* = 6) of 181-744 proteins explains 84% of the variance in the data (Figure 4A,B).

**Figure 4.**
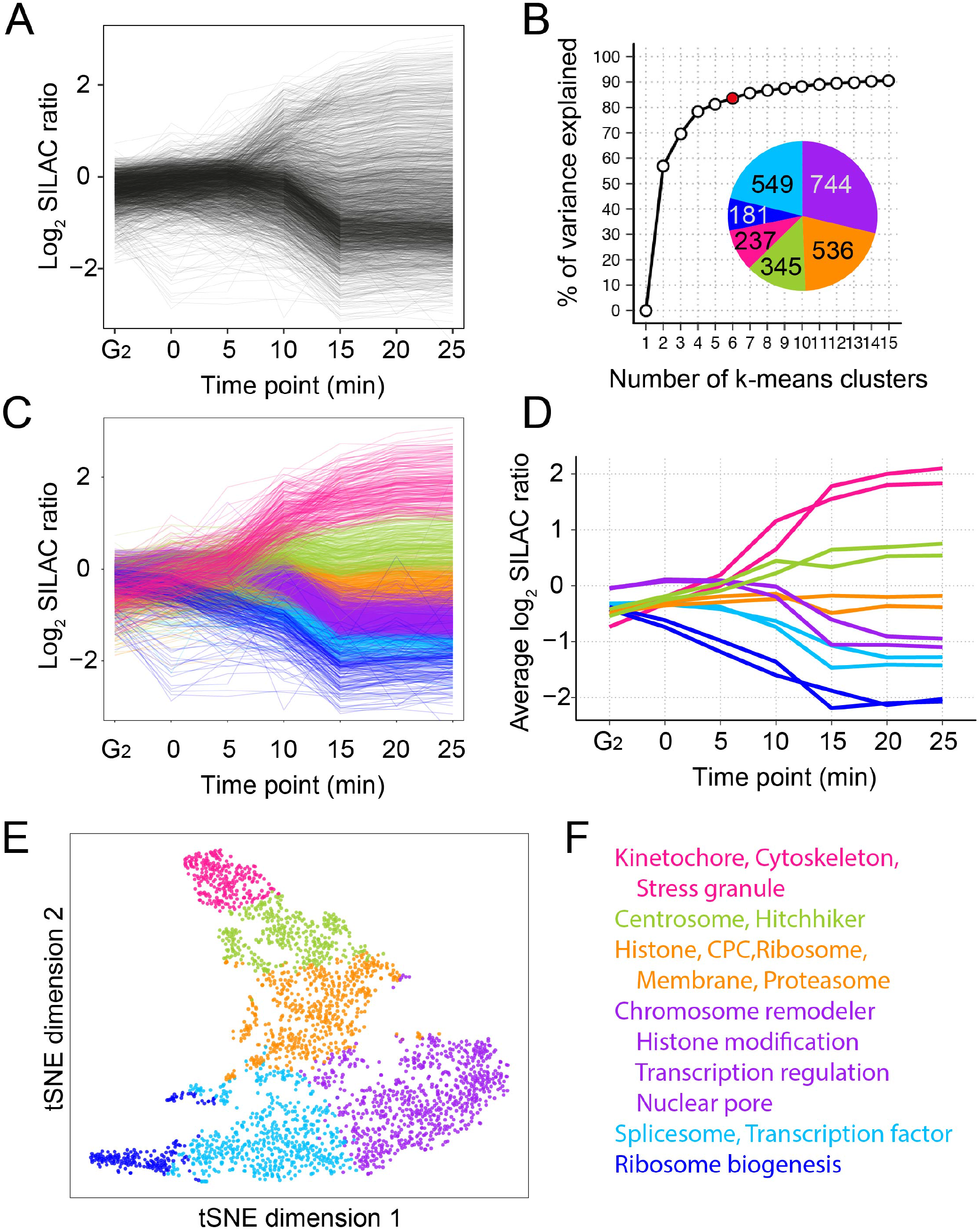
Over-all patterns of chromatin protein behavior during prophase. (A) Line plots showing the SILAC ratios of all proteins in the time course. Data from replicate A is shown. Reproduced from Figure 2A. (B) Percentage of variance explained as a function of the number of *k*-means clusters (red circle: k=6). Pie chart shows the number of proteins assigned to each of the six *k*-means clusters. This coloring for each cluster is used in all subsequent panels. (C) Line plots as in Panel A color-coded according to the 6 *k*-means clusters. (D) Line plots showing the median SILAC ratio of each cluster. Data for both replicates are shown. (E) T-distributed stochastic neighbor embedding (t-SNE) plot of the dataset in two dimensions. (F) Functional classification of representative proteins in each cluster.

Most chromatin proteins significantly increase or decrease their proximity to chromatin during prophase, however, about 20% remain relatively unaffected (Figure 4; orange cluster). This behavior was well reproduced between biological replicates (Figure 4D). For example, the chromatin association of 181 proteins decreases strongly immediately after mitotic entry (dark blue cluster). Others decreased only after 10 minutes and to a lesser extent (purple cluster). As expected from Figure 2, remodeling of the chromatin proteome was largely complete by the time of NEBD. None of the six groups of proteins showed further significant changes after 15-minutes (Figures 2, 4C,D).

We used t-distributed stochastic neighbor embedding (t-SNE) for an alternative visualization of this proteomic time course. t-SNE takes a dataset with many dimensions (e.g., two replicates, each with 7 ChEP time points) and reduces it to 2 dimensions (Van Der Maaten and Hinton, 2008). In the t-SNE plot (Figure 4E), each point corresponds to a single protein and the proximity between points reflects how similarly the proteins behave across the mitotic progression time course. The six groups of proteins identified by *k*-means clustering occupy distinct areas of the t-SNE plot, which resembles a “UK map”. Proteins along the “south coast” are decreased on chromatin as cells enter mitosis, with those in the west leaving first and those in the east leaving after NEBD. Proteins in the far north accumulate on chromatin during mitosis.

To assess changes in the absolute composition of mitotic chromatin, we estimated protein copy numbers and mass using intensity-based absolute quantification (iBAQ) (Figure S2) (Arike et al., 2012; Schwanhausser et al., 2011). In G_2_ cells proteins of the magenta cluster account for 1.2% of the ChEP-purified material in terms of copy numbers (Figure S2A, left) and 2.1% of the protein mass (Figure S2 B, left). However, these same proteins make up 8.9% (copy number) and 17.6% (mass) of the protein associated with mitotic chromosomes, the biggest enrichment seen for any cluster (Figure S2A,B, right). Of the other proteins, only the green cluster was significantly increased on mitotic chromosomes. All other clusters decreased (though note that the portion of the orange cluster corresponding to core histones, marked by a black arc, remained constant).

The different prophase behavior patterns are associated with proteins having distinct biological functions. Thus, each of the six *k*-means groups is significantly enriched for a specific set of Gene Ontology (GO) terms (Figure 4F, Table S1,S2) (Ashburner et al., 2000; Gene Ontology, 2021). For example, proteins that are quickly and strongly enriched on mitotic chromosomes (magenta cluster) belong to kinetochores, the cytoskeleton and stress granules. On the other hand, ribosome biogenesis factors (blue) are rapidly depleted from chromatin during earliest prophase.

### Hierarchical clustering reveals the behavior of specific protein groups

Groups defined by *k*-means clustering are large and functionally diverse and reveal little about behavior of specific functional groups of proteins. We therefore used hierarchical clustering for a more fine-grained analysis of the chromatin proteome.

The granularity of clustering analysis is adjusted by altering the height (h) of the cut of the dendrogram. In the analysis of Figure 5A, h = 1.7 assigned 2,592 proteins to 83 clusters: 38 clusters with 2 - 904 proteins and 45 with a single protein. Because we will use different values of h to reveal fine-grained features within certain clusters, we refer to cluster ‘X’ from this level of clustering as X_/83_. Four large clusters decrease their chromatin association during mitotic entry. In contrast, proteins that increase on the chromatin show a more complex behavior with eleven distinct clusters (Figure S2).

**Figure 5.**
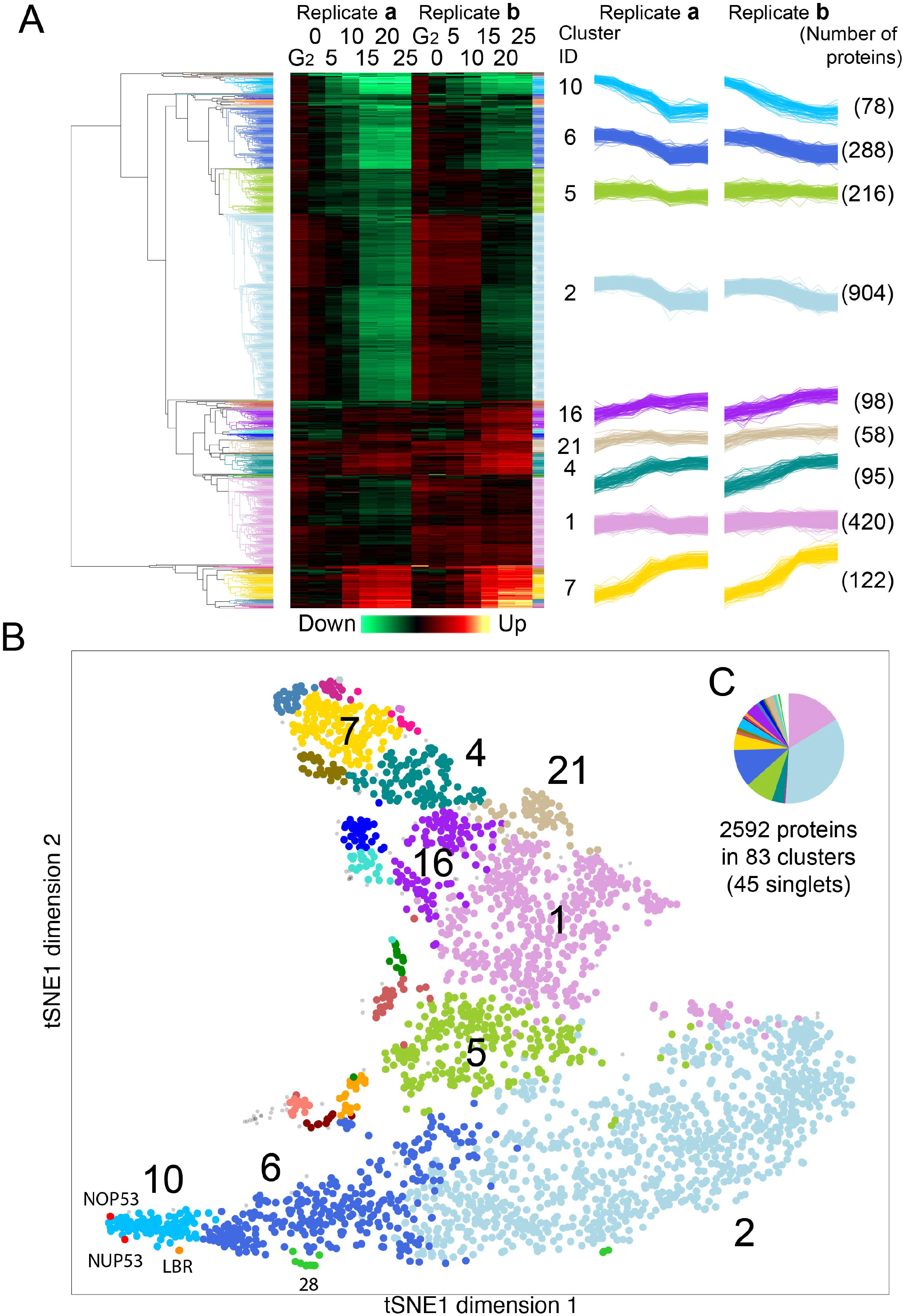
Hierarchical clustering of ChEP datasets. (A) Heatmap and line plots illustrate SILAC ratio of proteins from two replicates in time course experiments. Dendrogram shows hierarchical clustering. Each cluster is highlighted using a characteristic color throughout this figure. Line plots show kinetic behavior of proteins in the largest nine clusters. Cluster ID and number of members are given. (B) Illustration of hierarchical clustering results on a tSNE map. Cut tree height = 1.7. 23 clusters with 8 or more members and 2 singlets are colored, with the nine largest clusters annotated (see Figure 5A Suppl for full annotation). (C) Pie chart shows the number of proteins in 83 categories by hierarchical clustering.

General trends plotted for nine of the largest clusters highlight the reproducibility of the two biological replicates Figure 5A (right) and the average trend is shown for other selected clusters in Figure S3. Two major clusters, numbers 10_/83_ and 6_/83_, with 78 and 288 members, respectively, leave chromatin from the start of prophase (far south-west on the tSNE map). Levels of proteins of Cluster 10_/83_ in chromatin start to decline immediately upon release of cells from the G_2_ arrest. The proteins of Cluster 6_/83_ leave the chromatin later, after nuclear envelope permeability is compromised but with laminB1 showing rim localisation (Figure 3).

Clusters 1_/83_ and 2_/83_ dominate both G_2_ and mitotic chromatin. The 420 proteins of Cluster 1_/83_ account for 24% of the chromatin mass at G_2_ and 27% in mitosis. Cluster 1_/83_ is dominated by the core histones, which account for 11% of the calculated protein mass in both G_2_ and mitosis. Cluster 2_/83_, with 904 members (south-east on the tSNE map, light blue), is by far the most numerous cluster with 35% of the proteome (Figure 5C). Its members are depleted from chromatin starting after 10 min, coincident with nuclear envelope breakdown (NEBD) defined by lamin B1 fragmentation (Figure 3). These proteins include many “interphase chromatin proteins” (Kustatscher et al., 2014a; Kustatscher et al., 2014b) and account for 44% of the G_2_ chromatin mass (Figure S2D). Despite the substantial decrease in their chromatin-association after NEBD, they remain major components of mitotic chromosomes (27% of mitotic chromatin mass - Figure S2C,D).

Increasing the granularity of the analysis by setting h = 1 yielded 330 clusters (including 187 singlets). This divided the 1270 proteins of clusters 10_/83_, 6_/83_ and 2_/83_ (Figure 6A) into 10 subclusters with >10 members, plus numerous smaller clusters and singlets (Table 1 and Figure S4 A). Plots using this fine-grained analysis reveal that the tSNE map accurately reflects progressive trends early in mitosis (Figure 6B). From west to east across the south we observed discrete waves of protein exodus from chromatin. Further increasing the granularity (h = 0.6 - 964 clusters/680 singlets) can yield additional information about some closely related groups of proteins (Figure S4B; see next section), but the increased number of singlets limited its utility for functional insights.

**Figure 6.**
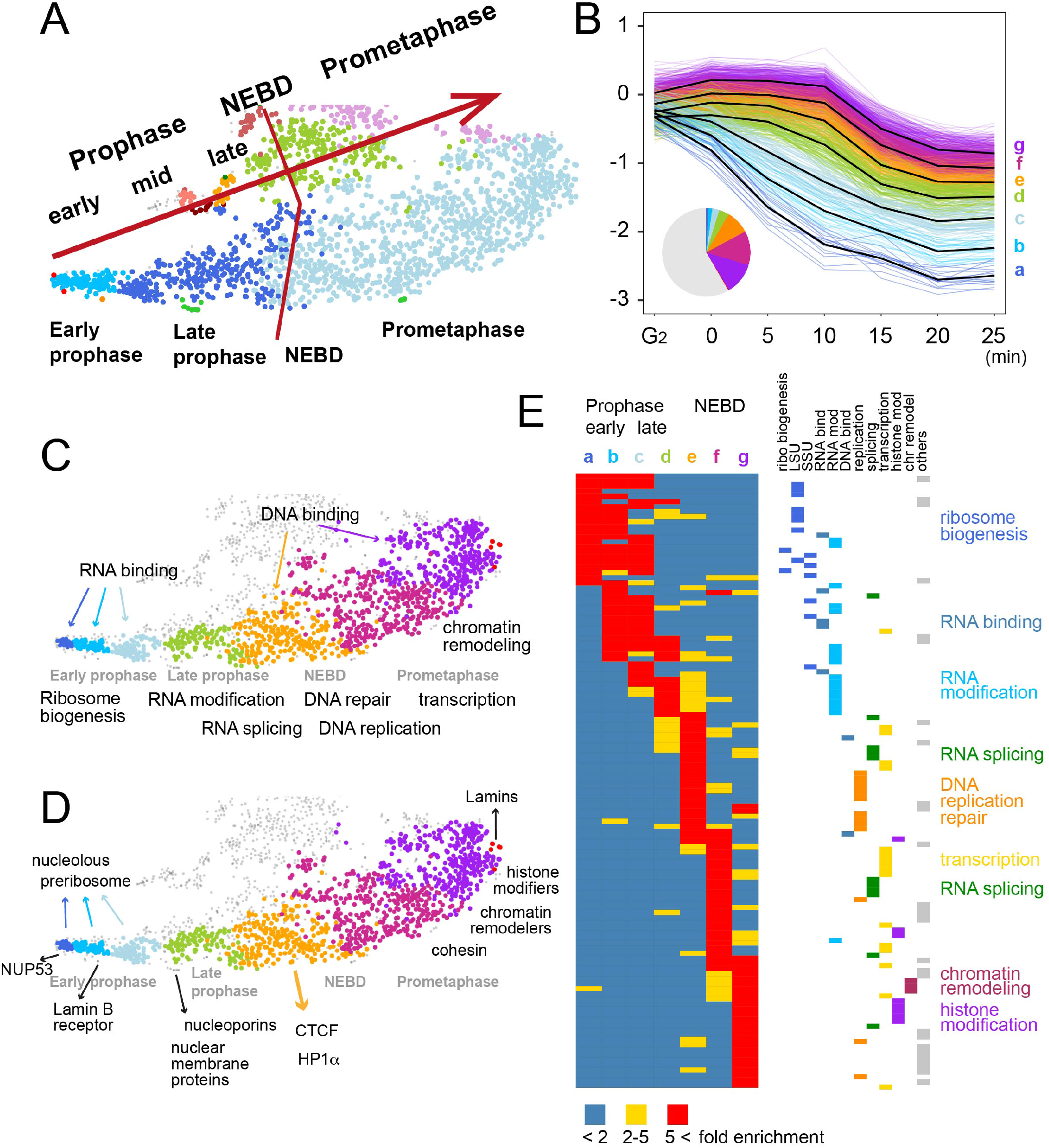
Interphase proteins leave chromatin sequentially in functional groups. (A) Correspondence of the tSNE map to events of mitotic progression clustered at tree height h = 1.7, as in Figure 5. Further cutting the tree height (h = 1.0) sub-divides clusters 10_/83_, 6_/83_ and 2_/83_ into 7 major subclusters, shown with new coloring (a-g) in panels B-D. (B) SILAC ratios of protein subclusters leaving chromatin. Black line: average for each subcluster. Pie chart: fraction of total protein IDs in each subcluster. (C) GO keywords for the tSNE distribution of Panel B subclusters. (D) Examples of specific proteins and functional classes. (E) GO ribbon table (left) showing subcluster matrix at h=1. Columns with color-coded labels show GO terms enriched in each subcluster. GO terms (rows) are color coded according to major categories. See Table 2 for details of individual GO terms.

### Very few proteins are unchanged on chromatin during mitotic entry

Cluster 1_/83_ contains 420 proteins whose levels change relatively little during mitotic entry. Within this cluster, 22 of the proteins (comprising subclusters 158_/964_ and 375_/964_) showed the least variation across the entire time course (see Figure S5D). In addition to the core histones and H2A.Z, these invariant proteins comprise an interesting group that might not have been predicted *a priori*. This group includes kinetochore proteins (CENP-C, CENP-I, Mad1 and KNL1), CPC members (Aurora B, INCENP, borealin), shugoshin, condensin II subunits CAP-H2 and CAP-G2, SMC5, SMC6 and PP2A B56γ.

### Reorganization of interphase chromatin during mitotic entry

Chromatin organization at the nuclear periphery is remodeled early in prophase, with most changes to the chromatin fiber organization elsewhere following only significantly later. HMGN1 and HMGA1 are among the first proteins to leave chromatin in Cluster 10_/83_. These DNA binding proteins regulate the higher-order structure of interphase chromatin (Catez et al., 2004; Postnikov and Bustin, 2016). However, other chromatin-organizing proteins leave the chromatin much later (after NEBD in Cluster 2_/83_). These include HP1α, cohesin, HMGBs, HMG20s, chromatin modifiers, transcription factors, mediator and integrator components (Figure 6).

In addition to regulating traffic between the nucleus and cytoplasm, nuclear pores also help regulate chromatin activity (Iglesias et al., 2020; Ptak and Wozniak, 2016; Van de Vosse et al., 2013). Nup53, a member of the inner nuclear pore ring and confirmed Cdk1 substrate (Linder et al., 2017), is one of the earliest proteins to move away from chromatin (Figures 5B, 6D). This is followed shortly thereafter by a cluster of four nucleoporins and four nuclear inner membrane proteins. The nucleoporins, NDC1, POM121C, NUP210, NUP210L, link the nuclear pore inner ring complex to the pore membrane (Kim et al., 2018; Mitchell et al., 2010). Whether changes in inner pore ring interactions with chromatin would influence pore barrier function is not clear, however these changes occur concomitant with weakening of the barrier between nucleus and cytoplasm. Other nucleoporins that can be crosslinked to chromatin leave at scattered times after Nup53 (Figure S5B).

Chromatin release from the nuclear envelope is essential for chromosome formation and segregation in mitosis (Champion et al., 2019). Consistent with this, another very early protein to leave mitotic chromatin is the inner nuclear membrane protein Lamin B receptor (LBR). LBR binds lamin B and heterochromatin protein HP1α (Ye and Worman, 1996) and may help target heterochromatin to the inner nuclear envelope. Cdk1 phosphorylation was reported to antagonize LBR binding to chromatin (Courvalin et al., 1992; Takano et al., 2004). LBR release from chromatin is followed shortly thereafter by other nuclear envelope transmembrane proteins, including LAP2 and MAN1 which have LEM domains that bind the chromatin tethering/crosslinking protein BAF (Dechat et al., 2000). Thus, HP1α-containing and BAF-containing heterochromatin may persist during the earliest stages of mitotic chromosome formation, but they are no longer tethered to the inner nuclear membrane.

Unexpectedly, the A and B-type lamins comprise one of the last subclusters of interphase chromatin-associated proteins to leave the chromatin.

### Nucleolar chromatin remodeling dominates early prophase

GO analysis reveals that proteins exit from chromatin in successive waves as cells progress through prophase. Figure 6C,D highlights selected individual proteins and functional classes. Surprisingly, earliest prophase is dominated by changes in the nucleolus (Figure 6E). The changes in chromatin association discussed below could reflect both disassembly of the chromatin and decreased assembly of new pre-ribosomal complexes.

The first cluster of proteins to leave the chromatin (cluster 10_/83_) is rich in nucleolar factors involved in RNA binding and ribosome biogenesis [Figure 6 B-E-a,b (blue)]. Cluster 6_/83_, which leaves slightly later, contains both nucleolar and non-nucleolar proteins [d (green) and e (orange)]. NOP53/PICT1 is the earliest of the nucleolar proteins to leave chromatin, already at t = 0. Nop53 is a key factor for pre-rRNA processing that targets rRNA to Mtr4 helicase associated with the exosome (Thoms et al., 2015). Mtr4 and the exosome remain in the chromatin until NEBD without Nop53, but pre-rRNA is presumably no longer targeted to exosomes. This likely interferes with processing and assembly of the 60S large ribosomal subunit (LSU). Indeed, proteins of the LSU processome are also depleted in chromatin starting at the 0-minute time point. Down-regulation of preribosome assembly is also driven by early removal of c-Myc, which coordinates ribosome production by transcriptional regulation of biogenesis factors (Destefanis et al., 2020; van Riggelen et al., 2010).

Interestingly, RNA polymerase I, several of its key transcription factors and its termination factor TTF1 (Evers et al., 1995) behave very differently from the pre-rRNA processing proteins, leaving the chromatin only after NEBD (10-15 minutes, Figure 7A,C). This suggests that pre-rRNA transcription may continue after rRNA processing ceases, potentially resulting in an accumulation of pre-rRNA, which ends up in the mitotic chromosome periphery compartment (MCPC) (Sirri et al., 2016).

**Figure 7.**
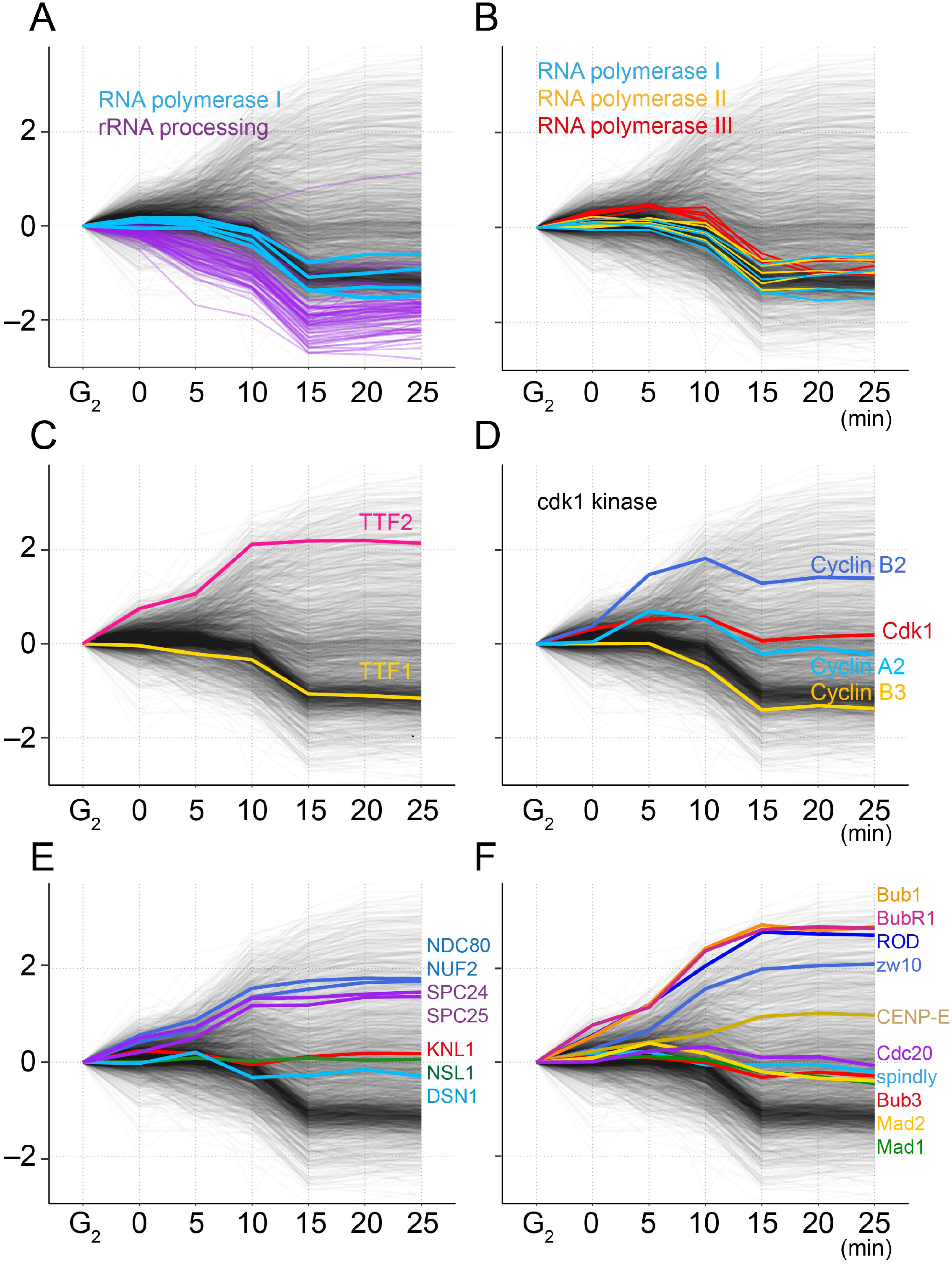
Behavior of selected proteins during mitotic entry. (A-F) Profile plots of proteins of interest, including ribosome biogenesis factors, RNA polymerases and termination factors, Cdk1 kinase and cyclins and kinetochore proteins, as indicated. Bulk proteins in the proteome are in grey.

Cluster 10_/83_ is enriched for proteins that interact with NPM1 and Surf6, two proteins that are important for nucleolar liquid-liquid phase separation (LLPS) (Figure S5A) (Ferrolino et al., 2018). Many nucleolar proteins have intrinsically disordered regions that are likely to participate in LLPS (Stenstrom et al., 2020) and many of these proteins end up in the MCPC, which forms much later in prometaphase and metaphase (Figures 6; S5A) (Sirri et al., 2016). Thus, association of the nucleolar phase with chromatin is reduced long before morphological changes are evident in the nucleolus by light or electron microscopy.

Proteins associated with mature ribosomes behave very differently from those involved in ribosome biogenesis (Figure S2A, S5A). They are released from the chromatin early, but begin to accumulate again later. This late accumulation may reflect the association of stress granule components with chromatin (Cluster 17_/83_ – see below).

Paradoxically, despite this mass exodus from chromatin, some proteins of Cluster 10_/83_ remain among the most abundant proteins associated with mitotic chromosomes.

### Interphase proteins leave chromatin in functional groups

Importantly, not all RNA processing factors are affected equivalently as cells enter prophase (Figure 6). As discussed above, the nucleolar chromatin and pre-rRNA processing machinery are disassembled in earliest prophase. In contrast, processes linked with RNAPII transcription are disassembled later. For example, factors involved in RNA splicing leave chromatin in late prophase, and many factors involved in DNA transcription, including histone modifiers and chromatin remodelers leave in the general exodus that follows nuclear envelope breakdown (e.g., Cluster 2_/83_).

RNAPII itself (and RNAPIII) are among the last of the proteins to leave the chromatin (Figure 7B). However, RNAPII termination factor TTFII increases on chromatin from the start of prophase, so RNAPII transcription may be shut off earlier than RNAPI transcription.

In order to look for finer patterns amongst the large number of proteins that leave chromatin following NEBD, we re-analyzed the data after setting h = 0.6. This divides the dataset into 964 units: 284 subclusters and 680 singlets. The eight largest subclusters are found within cluster 2_/83_ and comprise 525 proteins (54.8% of Cluster 2_/83_ - 20.3% of the entire dataset) (Figure S4B, Table S1). GO terms enriched among these clusters are relevant to DNA repair, DNA replication, chromatin remodeling, histone modifications, transcription initiation and some mRNA splicing (other splicing factors leave the chromatin earlier - Figure 6E for h=1). The various GO subclusters tend to leave the chromatin in waves. Importantly, as stated above, although the proteins in these subclusters reduce their abundance on chromatin they do not vanish from it entirely. They are still major components of mitotic chromosomes (Figure S2).

### Behavior of proteins that accumulate on mitotic chromatin is more diverse

Two protein clusters begin to associate with chromatin ahead of nuclear envelope breakdown (NEBD) (Figure S2B,C). Both are mostly composed of microtubule associated proteins, including several outer kinetochore proteins. Interestingly, the first (cluster 13_/83_) is the only cluster significantly enriched for Cdk1 substrates. Because most of the proteins in these two clusters are generally assumed to be cytoplasmic, their accumulation on chromatin well before NEBD is made possible by the early loss of nuclear envelope barrier function (Figure 3).

### Kinetochore proteins show a variety of behaviors

The kinetochore is an elaborate assembly of multi-protein complexes that assembles at centromeres to regulate mitotic progression and chromosome segregation (Hara and Fukagawa, 2018; Navarro and Cheeseman, 2021). The CCAN and Mis12 complex (e.g. NSL1, DSN1), which comprise the centromere-proximal portion of the kinetochore, show no change in association with chromatin from G_2_ through to prometaphase (Figure 7E, S5C). The CCAN components CENP-C and CENP-T recruit the tetrameric microtubule-binding NDC80 complex (NDC80-NUF2-SPC24-SPC25) (Gascoigne et al., 2011; Nishino et al., 2013; Rago et al., 2015), which is centrosomal during most of interphase, but moves into nuclei in early prophase (Hori et al., 2003). The NDC80 complex begins to accumulate on chromatin from time 0 in our samples. Its recruitment to chromatin is complete before NEBD, at 10 minutes (Figure 7E). Thus, kinetochores are presumably competent to capture cytoplasmic microtubules as soon as NEBD starts, and chromosomes are exposed to the cytoplasm.

Components of the MCC and RZZ spindle assembly checkpoint (SAC) complexes assemble in a stepwise fashion during mitotic entry (Figure 7F). The MCC components MAD1, MAD2 and CDC20 remain relatively invariant on chromatin throughout mitotic entry. The fourth MCC component, BubR1, is recruited later, starting in early prophase, in a cluster containing several other microtubule-associated proteins that is enriched for Cdk1 substrates (Cluster 13_/83_, mentioned just above).

Spindly, an adapter protein involved in RZZ recruitment, is stably associated with the chromatin from G_2_ onwards. In contrast, RZZ and dynactin, to which Spindly binds (Gama et al., 2017; Griffis et al., 2007), are recruited later in prophase. Accumulation of SAC components on chromatin continues even after NEBD starts.

The astrin/kinastrin complex (Dunsch et al., 2011), which stabilizes end-on microtubule attachments at kinetochores during metaphase (Conti et al., 2019) also associates with chromatin very early during prophase. Chromatin-associated astrin/kinastrin may help recruit stress granule components (Thedieck et al., 2013), many of which accumulate on chromatin after NEBD. It may also contribute to regulating separase activity at centromeres (Thein et al., 2007). Separase also undergoes a dramatic accumulation on chromatin during mitotic entry.

### Hitchhikers exhibit diverse behaviors

Hitchhikers differ from conventional contaminants (e.g., mitochondria). They are unlikely to function in chromosome formation or segregation but are physically associated with the chromosomes before cell lysis and apparently cannot be separated by conventional purification protocols (Ohta et al., 2010). Interestingly, some hitchhikers accumulate on chromatin, and may even plateau before lamina disassembly. For example, one cluster enriched with centrosomal proteins associates with chromatin very early - between G_2_ and 0 min then remains flat afterwards (Figure S3C - cluster 21_/83_). Accumulation of these cytoplasmic proteins on chromatin must either reflect the loss of nuclear envelope barrier function or be an artefact of the ChEP protocol.

Another cluster, number 7_/83_ with 122 proteins, increased significantly on chromatin over our time course (1% in G_2_ to 11% at 20 min). GO analysis reveals that this cluster is enriched for cytoplasmic proteins, including components of clathrin and COP-1-coated vesicles, mitochondria, cytoskeleton and endoplasmic reticulum (Tables S1; S2). Several other clusters that accumulate on chromatin are also significantly enriched for proteins that are more concentrated in ChEP fractions than on isolated chromosomes. Most members of these clusters are cytosolic, so these proteins are either hitchhikers or ChEP artefacts that are not true constituents of mitotic chromatin.

A late cluster of proteins to accumulate on chromatin is enriched in components of cytoplasmic stress granules (Figure S3C - cluster 17_/83_). Canonical stress granules are absent from mitotic cells (Ivanov et al., 2019; Riggs et al., 2020), possibly because polysomes disassemble during mitosis. Cessation of translation during mitosis may trigger an alternative response in which G3BP1, which is required to form the phase separating scaffold that drives stress granule formation (Sanders et al., 2020; Tourriere et al., 2003; Yang et al., 2020), associates with the condensing chromosomes and recruits other stress granule components.

## DISCUSSION

In this study we have mapped chromatin changes that accompany nuclear disassembly and mitotic chromosome formation using a proteomic approach based on crosslinking technology similar to that used in chromatin immunoprecipitation (ChIP). Our study defines a sequence of chromatin remodeling events, including release of large numbers of proteins from chromatin in successive waves accompanied and interleaved with the binding of cytoplasmic proteins to chromatin. These data are readily interrogated using a dedicated web app at https://mitoChEP.bio.ed.ac.uk.

### Cytoplasmic components access the chromatin early in prophase

Our ChEP analysis revealed that important interactions between cytoplasmic proteins and chromatin occur prior to nuclear envelope breakdown. Microscopy analyses confirmed that the nuclear-cytoplasmic barrier is lost within minutes of release from a G_2_ block, possibly starting with movement of Nup53 and inner pore ring components away from chromatin. Our results reveal the functional consequences of this precocious loss of nuclear envelope barrier function, which had been observed previously [see e.g. (Dultz et al., 2008; Lenart et al., 2003)], and may be driven by Cdk phosphorylation of nuclear pore components (Linder et al., 2017). For example, the completion of NDC80 complex association with chromatin prior to NEBD can explain the extremely rapid formation of bipolar attachments by chromosomes on the spindle after NEBD.

### Surprising order of chromatin remodeling during mitotic entry

Early prophase chromatin is characterised by an orderly transition as proteins leave chromatin in successive waves that form relatively tight clusters in our analysis. We expected that these early events might involve chromatin changes required to shape mitotic chromosomes. Indeed, HMGN1 and HMGA1 are two of the earliest proteins to leave chromatin. However most other interphase chromatin components, including cohesin and those involved in activities such as chromatin modification, remodelling, transcription and repair only change significantly after NEBD, long after prophase chromosome formation is complete.

Other early changes focus at the nuclear periphery, including chromatin release from nuclear pores and the nuclear membrane. Components of the nuclear pore have been implicated in interactions with ribosomal genes and heterochromatin (Iglesias et al., 2020; Ptak and Wozniak, 2016; Van de Vosse et al., 2013). Early release from the NPC may be linked to the other changes in the nucleolus that occur early in prophase (see below). Nuclear envelope proteins that reduce their chromatin interactions early include Lamin B receptor and several LEM-domain proteins. Apparently, chromatin rich in HP1α and BAF loosens its association with the inner nuclear membrane early in prophase before loss of HP1α (chicken BAF is not seen in our dataset). This chromatin release from the nuclear envelope is essential for chromosome formation and segregation in mitosis (Champion et al., 2019). Unexpectedly, the chromatin-associated populations of lamins A and B are among the very last proteins to detach from DNA in prometaphase.

The timing of protein release from chromatin is not simply proportional to the extent of direct Cdk1 phosphorylation. Thus, despite the presence of several known Cdk1 substrates in cluster 10_/83_ (e.g., NPM1, NPM3, NCL), Cdk1 substrates are not particularly enriched in this first large cluster to leave chromatin. Most Cdk1 substrates change their chromatin association later in prophase and prometaphase. This appears to correlate more with the behavior of cyclins than of Cdk1 itself (Figure 7D). Cdk1 itself shows relatively little variation in chromatin across our time course (cluster 1_/83_). Cyclin B2 accumulates on chromatin from G_2_ onward, peaking just before NEBD. Cyclin A2 and B3 levels fall in chromatin after NEBD. CDC25A/B leave chromatin in early prophase. Cyclin B1 is not yet identified in chicken.

### Unstressed nucleolar disassembly during mitosis

Surprisingly, GO analysis reveals that early prophase is dominated by changing associations of components involved in RNA-protein interactions including ribosome biogenesis, RNA modification and mRNA splicing. These early changes are particularly dramatic in the nucleolus and occur long before visible changes in nucleolar structure in late prophase/prometaphase. This makes sense, as nucleolar disassembly must occur during mitosis so that chromosomes carrying the ribosomal genes are free to segregate independently.

Inhibiting ribosome production during interphase triggers a nucleolar stress response sensed by Nop53/PICT1 (Sasaki et al., 2011) and 5S RNP (RPL5, RPL11 and 5S RNA) (Sloan et al., 2013; Weeks et al., 2019). The sensors respond by inhibiting MDM2, leading to activation of a p53 pathway culminating in cell cycle arrest or apoptosis (James et al., 2014; Yang et al., 2018). A second arm of this response involves c-Myc, a master regulator of ribosome biogenesis (Destefanis et al., 2020; van Riggelen et al., 2010). A common readout of the stress response is movement of NPM1 out of the nucleolus (Yang et al., 2018).

Interestingly, one of the earliest events of prophase is movement of NPM1 away from the chromatin. However, this process does not reflect nucleolar stress, as it would during interphase. Indeed, Nop53 is the first of the >2500 proteins to show a significant movement away from chromatin. Proteins found in the same cluster include c-Myc, RPF2 and RRS1. Removal of the latter two likely prevents incorporation of RPL5 and RPL11 into the LSU-processome (Zhang et al., 2007). We speculate that early removal of the sensors and c-Myc from chromatin provides a mechanism permitting nucleolar disassembly without activating the stress response during mitosis.

### Complex dynamics of the mitotic chromosome periphery compartment (MCPC)

The earliest protein clusters depleted from chromatin following the release from G_2_ are highly enriched in nucleolar components (clusters 10_/83_ and 6_/83_). Many of these components accumulate on the surface of mitotic chromosomes in the MCPC, but this association only occurs during prometaphase or even later in mitosis (Sirri et al., 2016). Indeed, that still-mysterious layer coating the chromosomes is apparently composed largely of nucleolar and pre-ribosomal proteins and RNAs (Booth et al., 2014; Stenstrom et al., 2020). The early release of these proteins from chromatin poses an interesting conundrum. The MCPC absolutely requires Ki-67 for its formation (Booth et al., 2014; Cuylen et al., 2016; Stenstrom et al., 2020). However, Ki-67 leaves chromatin much later than the other nucleolar MCPC components.

Many of the proteins of clusters 10_/83_ and 6_/83_ associate with NPM1, which together with Surf6 drives liquid-liquid phase separation (LLPS) during nucleolar formation. It has been speculated that the MCPC represents a separated phase coating the chromosome surface. Indeed, in the absence of Ki-67 MCPC proteins form what appear to be large phase condensates in the mitotic cytoplasm (Booth et al., 2014; Hayashi et al., 2017). The status of these proteins between earliest prophase, when they begin to move away from chromatin together with NPM1 and Surf6, and late prophase/prometaphase when the MCPC begins to form remains an interesting question for future research.

### Stress granule components associate with mitotic chromatin

Stress granules were among the first cellular structures shown to form as a result of LLPS (Molliex et al., 2015). They form when, in response to various stresses, translation initiation is inhibited and the ribosomes of polysomes “run off” the mRNA, leaving behind circular RNPs associated with 43S preinitiation complexes (Protter and Parker, 2016; Riggs et al., 2020). Canonical stress granules do not form during mitosis, when polysomes are absent. However, it is not clear why they do not form when translation is suppressed during mitotic entry. Interestingly, stress granule components comprise one of the clusters with the greatest enrichment on mitotic chromatin following NEBD. Whether stress granules themselves associate with mitotic chromosomes, or whether their components associate with the chromosomes separately remains to be determined. Our preliminary experiments have revealed over 18,000 mRNAs associated with isolated HeLa mitotic chromosomes. However further experiments are required to determine whether these mRNAs are present in stress granules and whether they contribute to formation of the MCPC.

### Perspectives

This first map of chromatin transactions during mitotic entry has revealed several surprises. Changes in RNP associations with chromatin, particularly in the nucleolus, occur long before most changes of canonical chromatin components. Furthermore, significant remodeling of chromatin by cytoplasmic proteins occurs in early prophase long before conventional NEBD. These and other aspects of our map can be explored interactively using a dedicated app at https://mitochep.bio.ed.ac.uk.

## ACKNOWLEDGEMENTS

We thank Natalia Kochanova and Shaun Webb for helping to refine and deploy the Shiny App and Lucy Remnant, Bram Prevo, Fernanda Cisneros-Soberanis, Caitlin Reid Jeyaprakash Arulanandam and Natalia Kochanova for comments on the MS. This work is funded by Wellcome grants: 107022 & 221044 to WCE and 203149 to the WCB. GK is funded by an MRC Career Development Fellowship (MR/T03050X/1).

## AUTHOR CONTRIBUTIONS

Experiments: IS, CS, KS. Data interpretation: IS, GK and JR. MS preparation: IS, GK and WCE.

## DECLARATION OF INTERESTS

The authors declare no competing interests.

## STAR METHODS

### Cell culture and medium

The chicken lymphoma B cell line DT40 with CDK1^as^ allele (Gibcus et al., 2018; Samejima et al., 2018) was grown in RPMI1640 medium supplemented with 100 μg/ml ^13^C_6_ ^15^N_2_-L-lysine:2 HCl, 30 μg/ml ^13^C_6_ ^15^N_4_-L-arginine:HCl, 10% dialyzed FBS (mol wt cut-off, 10,000) and 1% penicillin/streptomycin.

GFP-NLS Halo-LaminB1 cells and control cells for spike-in were grown in RPMI1640 medium supplemented with complete FBS. Cells were grown at 39°C, 5% CO_2._

### Synchronization of CDK1as cells

Cells were grown to 1×10^6^ cells/ ml. 1NM-PP1 was added to 2 μM and further incubated for 10 or 13 hours. The G_2_ arrested cells were washed three times with RPMI medium supplemented with 56 mM PIPES pH = 7.0 (RPMI-PIPES). Washed cells were resuspended in RPMI-PIPES at cell density of 1×10^6^ cells/ ml, aliquoted and incubated for a set time.

### Construction of 3xGFP-NLS plasmid

3Xsuperfolding GFP from addgene-plasmid-75385 (digested with BamHI/XhoI) and double strand oligos encoding BP-NLS (*tcga*gaagcgcgtaaccgcagcgggcatcacgcatccaaagaaaaagcggaaagtgtaag*ggcc* and cttacactttccgctttttctttggatgcgtgatgcccgctgcggttacgcgcttc) were ligated into pcDNA 3 (digested with BamHI/ApaI).

### Construction of DT40 Halo-lamin B1 cell line

A Hygromycin-resistant-ORF_P2A_Halo tag was inserted at the N-terminus of the lamin B1 gene using CRISPR/Cas9 gene editing technology. The guide RNA targeting Lamin B1 (TCCCCTACCATCACGTCACG) and Cas9 expressing plasmid (pX330) plus an N-terminus Halo tag knock-in construct (Hygromycin resistance gene-P2A-Halo) were co-transfected into wild type CDK1^as^ chicken DT40 cells using the NEON transfection system. After 48h, cells were transferred to 6 × 96-well plates in selective media (hygromycin 0.6 mg/ ml from Thermofisher).

In order to obtain cell lines expressing 3xGFP-NLS, Halo-lamin B1 knock-in cells were transfected with plasmid expressing 3xGFP-NLS by electroporation in a GenePulser (Bio-Rad). After 24 h in selective media (G418), GFP-positive clones were screened by flow cytometry and microscopy analysis.

### Mass spectrometry

Cells were fixed with 1% formaldehyde for 10 minutes. To inactivate the formaldehyde, 1/20 volume of 2.5 M glycine was added and incubated for 5 minutes before harvesting cells. The fixed cells were washed with TBS (50 mM Tris pH 7.5, 150 mM NaCl), and snap frozen in liquid nitrogen for storage at −80°C. Once thawed on ice, heavy and light labelled cells were mixed and processed according to the ChEP protocol (Kustatscher et al., 2014a; Kustatscher et al., 2014b). The DNA content of the chromatin fractions was measured using a Qubit with HS DNA QuantIT (Thermo Fisher Scientific) according to the manufacturer’s instructions.

ChEP chromatin was processed for mass spectrometry by in-gel trypsin digest. The detailed procedure is described in (Samejima and Earnshaw, 2018). In brief, tryptic peptides were fractionated by performing strong cation exchange chromatography, using a PolySULFOETHYL A (Poly-LC) column (Hichrom, UK). Mobile phase A consisted of 5mM KH_2_PO_4_, 10% acetonitrile at pH 3; mobile phase B was 5 mM KH_2_PO_4_, 1 M KCl, and 10% acetonitrile, pH 3. The peptides were fractionated using the following gradient: 0%–60% buffer B in 18 min, then to 70% in 2 min, and then to 0% in 6 min. The flow rate was constant at 200 μl/min. Fractions were collected at 1-min time slices. Fractionated samples were combined into six fractions. The peptide samples were desalted on C18 stage tips as described before (Rappsilber et al., 2003).

Mass spectrometry analyses were performed on a Q Exactive mass spectrometer (Thermo Fisher Scientific), coupled on-line to a 50 cm Easy-Spray HPLC column ES803 (Thermo Fisher Scientific), which was assembled on an Easy-Spray source and operated constantly at 50°C. Mobile phase A consisted of 0.1% formic acid, while mobile phase B consisted of 80% acetonitrile and 0.1% formic acid. Peptides were loaded onto the column at a flow rate of 0.3 μl min^−1^ and eluted at a flow rate of 0.25 μl min^−1^ according to the following gradient: 2 to 40% buffer B in 180 min, then to 95% in 11 min (total run time of 220min). Survey scans were performed at 70,000 resolution (scan range 350-1400 m/z) with an ion target of 1.0e6 and injection time of 20ms. MS2 was performed with an ion target of 5.0E4, injection time of 60ms and HCD fragmentation with normalized collision energy of 27 (Olsen et al., 2007). The isolation window in the quadrupole was set at 2.0 Thomson. Only ions with charge between 2 and 7 were selected for MS2.

All mass spectrometry raw files have been deposited to the ProteomeXchange Consortium (http://proteomecentral.proteomexchange.org) via the PRIDE partner repository with the dataset identifier PXD026385 (Reviewer account details: Username, Password: kz1F3sRu). MaxQuant version 1.6.7.10 (Cox and Mann, 2008) was used to process the raw files and peptide searches were conducted against the chicken reference proteome set of UniProt database (downloaded on April 2, 2020) with additional sequences from our in-house database of chicken proteins, using the Andromeda search engine (Cox et al., 2011).

### Data Analysis

Statistical analysis was performed with R (R_Core_Team, 2021) and Perseus (Tyanova et al., 2016). SILAC ratios reported by MaxQuant were log2-transformed and normalised such that the average log2 ratio of the four core histones was zero at each time point. Proteins detected in half or less of the 14 analysed samples (two replicates of seven time points) were discarded from the analysis. 2,592 of the remaining 3,500 proteins were detected in all 14 samples, and these proteins were used for statistical analyses that required complete data matrices (PCA, t-SNE and clustering).

The Rtsne package for R (Krijthe, 2015) was used to visualize the data by t-Distributed Stochastic Neighbor Embedding (t-SNE) (Van Der Maaten and Hinton, 2008). The theta parameter was set to zero to calculate the exact embedding. The perplexity parameter was set to 50, up from the default of 30, to account for the large size of the dataset.

We grouped proteins by *k*-means clustering. This divides a dataset into *k* groups based on how similar the behavior of each individual is to the mean behavior of its corresponding group across the time course. The base R function was used for *k*-means clustering, using the default algorithm by Hartigan and Wong (Hartigan and Wong, 1979).

Similarly, hierarchical clustering was performed using base R functions at standard settings (Euclidean distance and “average” agglomeration method). To call clusters at different levels of “granularity”, the clustering tree was cut at three different heights h (h = 1.7 for coarse clusters, h = 1.0 for medium clusters and h = 0.6 for fine-grained clusters).

Gene Ontology (GO) annotations for chicken were downloaded from the EBI GO Annotation Database (https://www.ebi.ac.uk/GOA/). The topGO R package (Alexa et al., 2006) was used to identify GO terms enriched in various clusters. Rather than the whole chicken proteome, only proteins that were included in the cluster analysis and had GO annotations were used as the gene ‘universe’ or background for the topGO analysis. Enrichment of GO terms in clusters was tested considering GO graph structure and using a Fisher’s exact test.

The web app that makes our results available as an interactive online resource at https://mitochep.bio.ed.ac.uk was created using R Shiny (Chang et al., 2021). The R code required to run the app can be accessed at: https://github.com/kustatscher-lab/mitoChEP-Shiny-App.

### Microscopy

Cells were fixed with 4% formaldehyde then washed with TBS (50 mM Tris pH 7.5, 150 mM NaCl) before spreading on a Polysine-coated slide. Attached cells were incubated with JF549 halo ligand (Chong et al., 2018) (a kind gift of Dr Luke Davis, Janelia Farm) followed by Hoechst 33542 (Invitrogen).

Fluorescent microscopy images were captured and processed using a legacy DeltaVison microscope system with SoftWorx software (Applied Precision Inc, lmage Solutions UK Ltd).

## SUPPLEMENTARY INFORMATION

### SUPPLEMENTARY FIGURE LEGENDS

**Figure S1. Comparison of replicate A and replicate B at each time point.** The Pearson Correlation Coefficient (PCC) is indicated.

**Figure S2 Suppl. Relative composition of the chromatin proteome at G_2_ and in mitosis (20 min).**

(A, B) Proteins grouped by *k*-means clustering (see Figure 4)

(A) Distribution of protein copy numbers (iBAQ algorithm) in the various *k*-means clusters.

(B) Calculation of the total protein mass in the various *k*-means clusters.

(C, D) Proteins grouped by hierarchical clustering (h = 1.7 - see Figure 5)

(C) Distribution of protein copy numbers (iBAQ algorithm) in the various hierarchical clusters.

(D) Calculation of the total protein mass in the various hierarchical clusters.

The black arc with asterisks indicates the contribution of the core histones (Histone H2A, Histone H2B, Histone H3, Histone H4).

**Figure S3. Line plots of average SILAC ratio of major clusters from selected hierarchical clusters (h=1.7).** Results from two replicates are shown in left and right columns, respectively.

**Figure S4. The largest protein clusters remaining after increasing the stringency of cut-off (decreasing the cut tree height, h) in hierarchical clustering.**

(A) At h = 1 the 16 largest clusters_/330_ comprise 1661 proteins

(B) At h = 0.6, 8 of the 9 largest clusters_/964_ derive from Cluster 2_/83_.

**Figure S5. Behavior of selected protein groups shown by mapping on the tSNE map and hierarchical clustering.**

(A) Diversity in kinetic profiles of nucleolar proteins identified using various algorithms illustrated on the tSNE map (h = 1.7 and cluster colors as in Figure 5). The embedded pie charts show the number of proteins in each group together with their cluster affiliation at h = 1.7.

(B) Nucleoporins do not dissociate from chromatin in a single cluster, but are spread across the tSNE map.

(C) Cdk1 substrates are spread across the tSNE map (left). Kinetics profile of 3 mitotic kinases (right). Plk1 shows a distribution very like Cdk1, whereas Aurora B is one of the small group of proteins that remain invariant on chromatin during mitotic entry.

(D) (left) Distribution of kinetic profiles of selected centromere / kinetochore proteins. Many of the proteins show relatively small changes in mitosis, but a number of them are significantly increased during mitotic entry (“north” in the tSNE map). (right) A line plot shows the changing levels of chromatin association for these proteins.

(E) The two subclusters derived from Cluster 1_/83_ containing the 22 invariant proteins that include the CCAN components CENP-C and CENP-T.

### SUPPLEMENTARY TABLES

**Supplementary Table 1:** List of proteins in ChEP samples, annotated with SILAC ratio, Uniprot database ID, *k*-means and heirarchical cluster names and GO terms.

**Supplementary Table 2:** Results of GO enrichment analysis for each cluster and detailed list of GO terms corresponding to the Y axis in Figure 6F.

## Notes

### Competing Interest Statement

The authors have declared no competing interest.

