## Supplementary figures and images for "Mapping the invisible chromatin transactions of prophase chromosome remodelling"

### Supplementary Figures S1 - S5

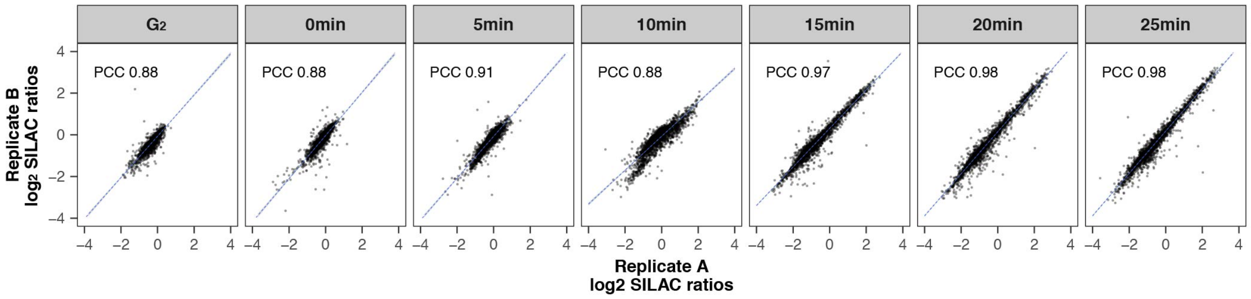

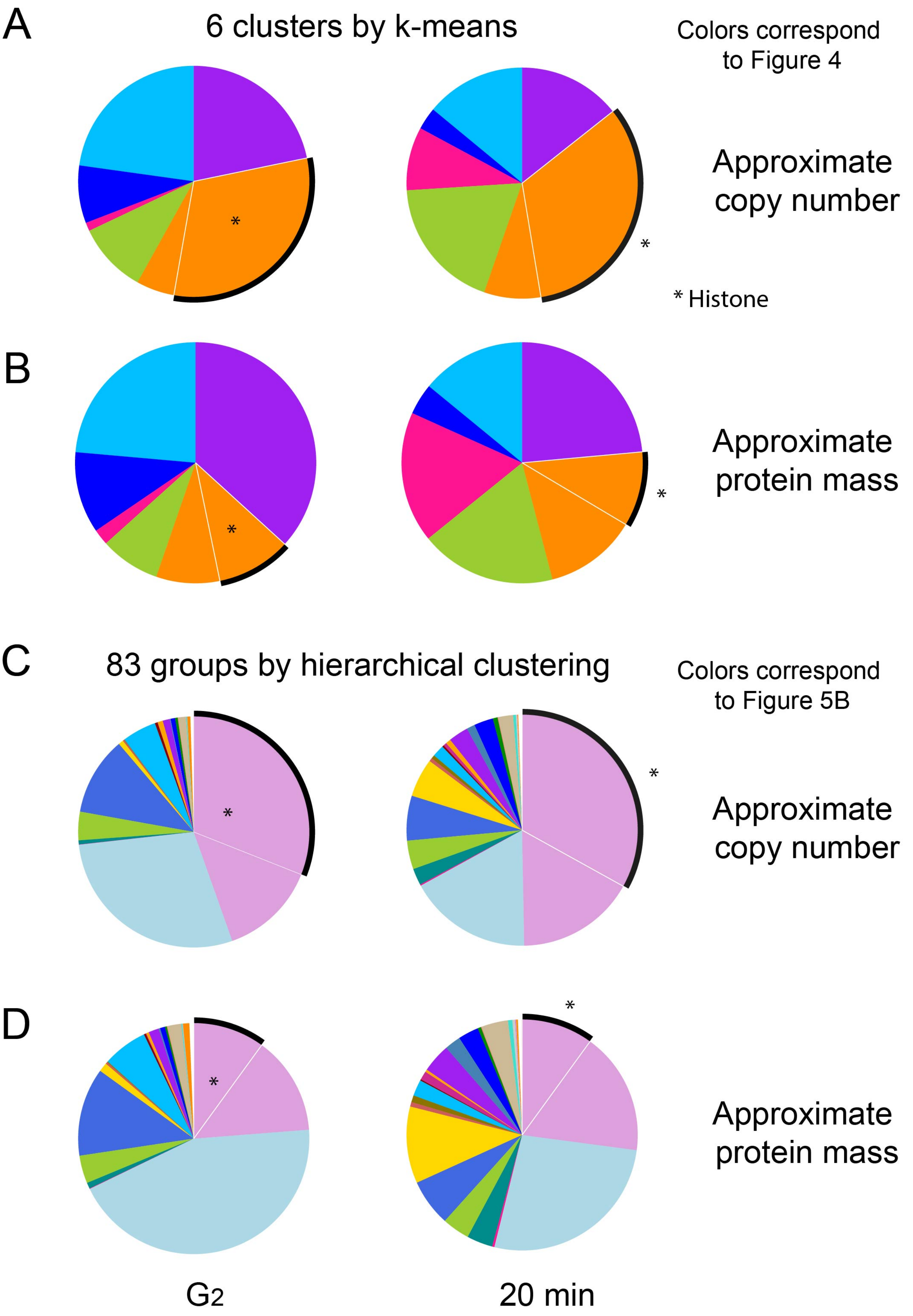

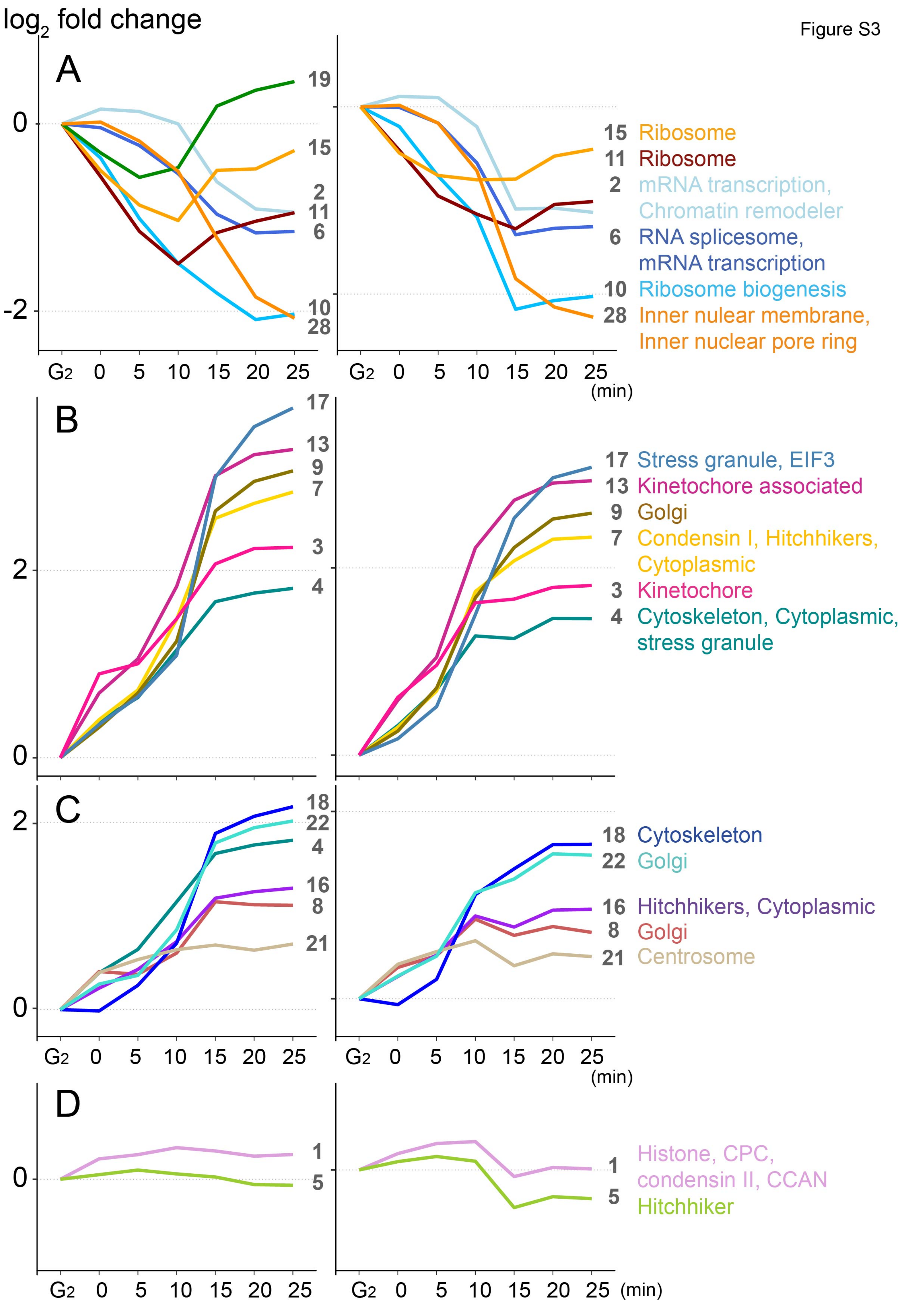

**A** 16 clusters with >30 members at h=1

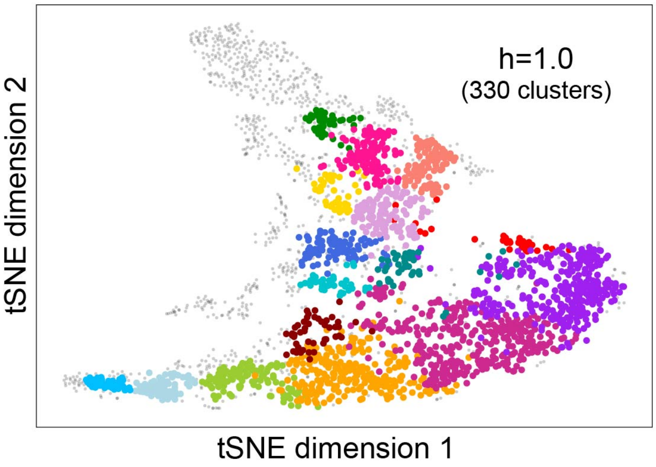


**B** Separation of Cluster 2/83 into 8 sub-clusters

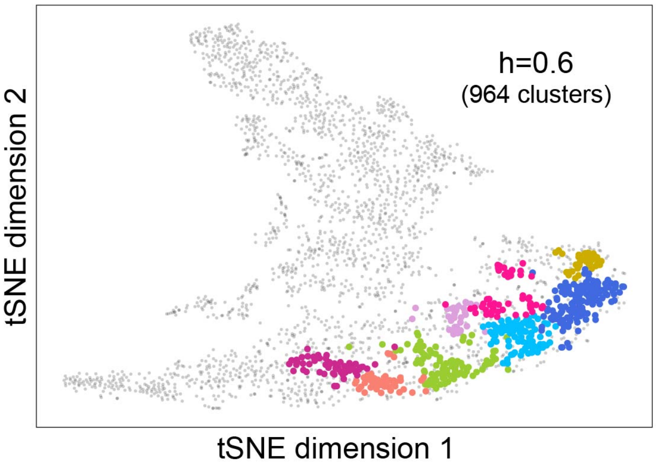

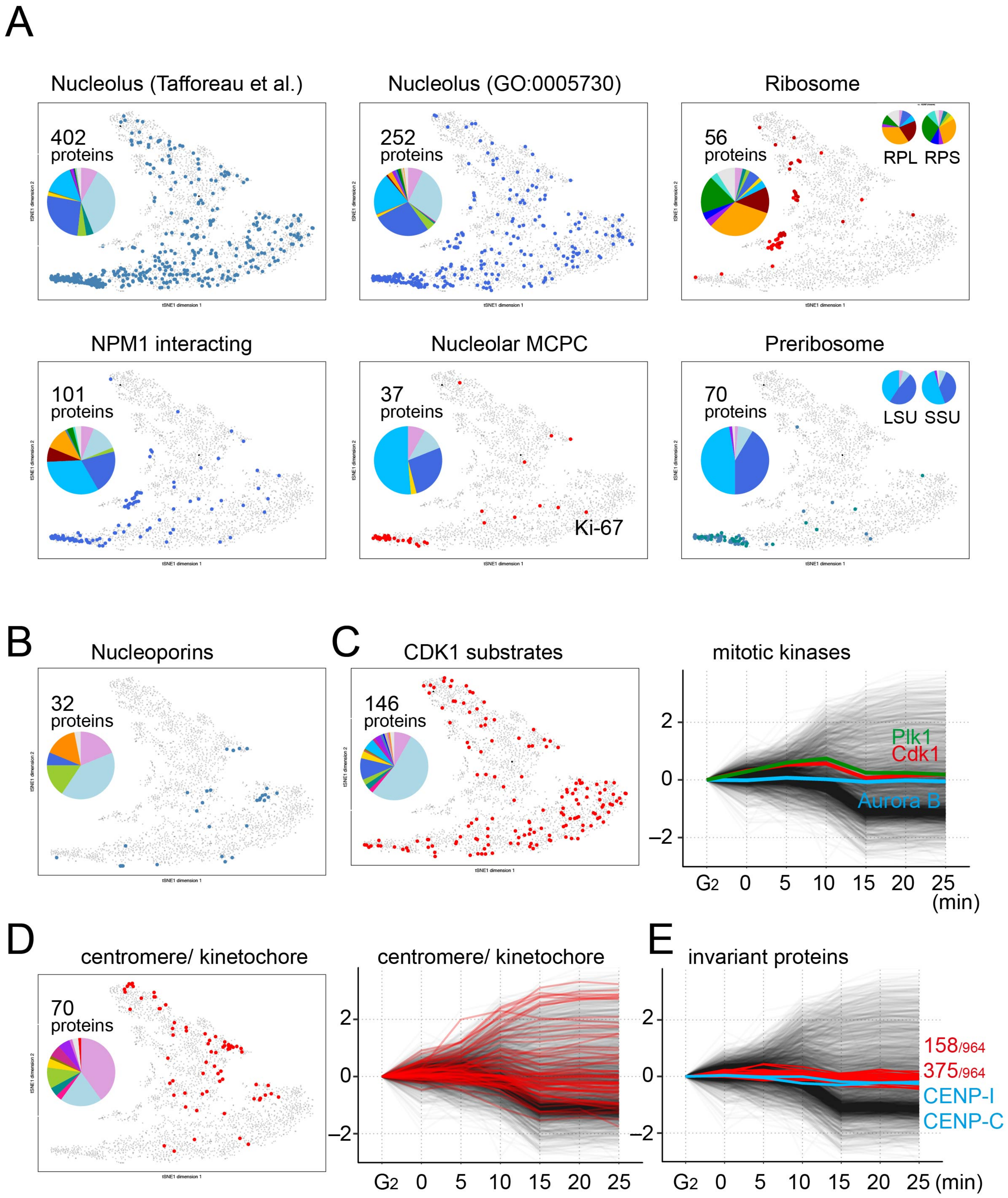
